# RECAS9: Recombining wild species introgression via mitotic gene editing in barley

**DOI:** 10.1101/2020.01.07.897280

**Authors:** Shelly Lazar, Manas Ranjan Prusty, Khaled Bishara, Amir Sherman, Eyal Fridman

## Abstract

Genetic loci underlying variation in traits with agronomic importance or genetic risk factors in human diseases have been identified by linkage analysis and genome-wide association studies. However, narrowing down the mapping to the individual causal genes and variations within these is much more challenging, and so is the ability to break linkage drag between beneficial and unfavourable loci in crop breeding. We developed RECAS9 as a transgene-free approach for precisely targeting recombination events by delivering CRISPR/Cas9 ribonucleotide protein (RNP) complex into heterozygous mitotic cells for the barley (*Hordeum vulgare*) *Heat3.1* locus. A wild species (*H. spontaneum*) introgression in this region carries the agronomical unfavourable tough *rachis* phenotype (non-*brittle*) allele linked with a circadian clock accelerating QTL near GIGANTEA gene. We delivered RNP, which was targeted between two single nucleotide polymorphism (SNPs), to mitotic calli cells by particle bombardment. We estimated recombination events by next generation sequencing (NGS) and droplet digital PCR (ddPCR). While NGS analysis grieved from confounding effects of PCR recombination, ddPCR analysis allowed us to associate RNP treatment on heterozygous individuals with significant increase of homologous directed repair (HDR) between cultivated and wild alleles, with recombination rate ranging between zero to 57%. These results show for the first time in plants a directed and transgene free mitotic recombination driven by Cas9 RNP, and provide a starting point for precise breeding and fine scale mapping of beneficial alleles from crop wild relatives.

## Introduction

This relationship between genetic polymorphism within a species and the phenotypic differences observed between individuals is critical for Identifying and determining the causal genes underlying variation in traits (Korte and Farlow, 2013). Quantitative trait loci (QTL) from crop wild relatives (CWR) are arguably the most valuable source for novel alleles conferring adaptation of modern crops to changing environments, owing to the harsh environment they inhibit at (Zamir, 2001). Nevertheless, only few CWR-QTL for agricultural traits were cloned and dissected to the molecular level (Salvi and Tuberosa, 2005), and such map-based-cloning efforts required screening thousands of plants to identify an utilize recombinants for the mapping (Frary et al., 2000; Fridman et al., 2000; Uauy et al., 2006) (Mao et al., 2010). This proportion of cloned genes out of the total mapped QTL for yield related traits is minute and recall for more efficient methodologies to achieve this in a reasonable and cost-effective way. Genome wide association studies (GWAS) are often complementary to QTL mapping and, when conducted together, they are supposed to lessen each other limitations and maximize mapping resolution (Brachi et al., 2010). GWAS increases resolution of the mapping due to long history of recombination within the gene pool. However, it still suffer from under consideration of rare alleles, which then lead to the phenomenon of “missing heritability” often observed where most of the phenotypic variation could not be explained by the genetic variants considered in the genetic model (Bloom et al., 2019). One partial solution to that was given in plants, where genome scans evolved to the point where large multi-parent populations attempt to bridge the advantages of QTL mapping and GWAS, e.g, nested-association-mapping (McMullen et al., 2009)(Maurer et al., 2015). However, combining the GWAS and QTL linkage approaches is still insufficient to map the individual gene and especially the variation within the gene of interest. For example, the wild allele of the *HsDry2.2* gene showed a significant grain yield increase in in water limitation (WL) as compared to no-effect under well water (WW) treatment (Merchuk-Ovnat et al., 2018). Nevertheless, the location of the QTL is delimited to 5.5cM on chromosome 2, which corresponds to app. 400Mbp on the physical map (Merchuk-Ovnat et al., 2018). Therefore, generating recombination every 100Kbp for finer mapping will require at least 4000 F2 plants, assuming recombination landscape is uniform, which of course is not.

In fact, recombination rates are highly variable between species, populations, individuals, sexes, chromosomes, and chromosomal regions (Stapley et al., 2017;Jensen-Seaman et al., 2004;Serra et al., 2018). In meiosis, homologous chromosomes pair and recombine during the prophase step of the first meiotic division, which can result in a reciprocal exchange, termed crossover (Mercier et al., 2015). Due to the significant roles of homologous recombination (HR) in meiosis, Fernandes et al., 2018 acted to enhance HR by abolishing key elements FANCM, RECQ4, FIGL1 genes which are responsible for meiotic crossovers reduction. The greatest effect was observed when combining *recq4* and *figl1* mutations, which increased the hybrid genetic map length from 389 to 3,037 cM. This corresponds to an unprecedented 7.8-fold increase in crossover frequency. Similarly, (Serra et al., 2018) simultaneously increased recombination via elevating E3 ligase gene *HEI10* (promote HR) and introduced mutations to helicase genes *RECQ4A* and *RECQ4B* (suppressors of HR). In mitosis homologous recombination or homology directed repair (HDR) is involved in the maintenance of somatic cell genomic stability by repairing DNA lesions or strand breaks. It also changes with response to exogenous stresses such as radiation, chemical exposure and replication error that induce DNA double-strand breaks (Amunugama and Fishel, 2012). Recently, studies attempted to increase recombination in mitosis via this cell repair mechanism by disrupting the non-homologous end joining (NHEJ) pathway (Verbeke et al., 2013; Kretzschmar et al., 2013; Jang et al., 2018). For example, attempts to modulate HR included the use of inhibitors for the NHEJ pathway like SCR7, a DNA ligase IV inhibitor, which was first identified as an anti-cancer compound, and considered as a potential NHEJ inhibitor (Ma et al., 2016). Chu et al., 2015 increase the efficiency of HDR 4–5-fold by suppressing NHEJ key molecules such as KU70, KU80 or DNA ligase IV by gene silencing, the ligase IV inhibitor SCR7 or the co-expression of adenovirus 4 E1B55K and E4orf6 proteins. In addition, Kwon et al., 2012 demonstrated that overexpression of OsRecQl4 and/or OsExo1, two genes encoding proteins involved in the processing of DSB, enhanced break HR. Vis-à-vis the ability to achieve high-resolution QTL mapping, these HDR manipulations provide major advancement, yet they work at a genome-wide rather than as a more directed approach to pin-point causal DNA variation.

Notably, the process in which nucleotide sequences are exchanged between two similar or identical molecules of DNA may occur either in meiosis or mitosis cell division. Recent utilization of CRISPR/Cas9 gene editing for mediating recombination in yeast and plants indicate that this process could be directed in mitosis using specific targeting gRNA molecules. In yeast, directing double strand breaks (DSB) using the CRISPR/Cas9 allowed the generation of a mapping panels with targeted recombination events spaced along a yeast chromosome arm. This allowed the fine mapping of manganese sensitivity to a single polymorphism in the transporter Pmr1(Sadhu et al., 2016).In tomatoes, (Filler Hayut et al., 2017) demonstrated targeted recombination between homologous chromosomes upon somatic induction of DNA double-strand breaks (DSBs) via CRISPR-Cas9. Targeted DSB among two alleles carrying different mutations in the PHYTOENE SYNTHASE (PSY1) gene resulted in yellow fruits with wild type red sectors forming via HR-mediated DSB repair. Both these studies demonstrate the power and potential of CRISPR/Cas9 system to accelerate and target recombination and trait mapping. Nevertheless, they rely on stable transformation of yeast or plants, and are less modular to include any locus of interest, mainly one harboring introgression with QTL for complex traits.

In this study, we initiated RECAS9, a transgene-free delivery of ribonucleotide proteins (RNP) CAS9/sgRNA complex to mitotic cells for generation of DSB, and acceleration of specific recombination between wild and cultivated homologous chromosomes. This is conducted on *Hordeum spontaneum Heat3.1*, a wild barley introgression that affect the circadian clock and reproductive output under heat. Besides, on the same introgression are closely linked and complementary genes, Btr1 and Btr2, which control *rachis brittleness* in barley(Pourkheirandish et al., 2015). Recessive mutations in any of these two genes turn the fragile *rachis* (*brittle*) into a tough *rachis* phenotype (non-*brittle*) (Fernández-Calleja et al., 2019) Therefore, linkage drag between the three loci from wild barley deemed maladaptive for agricultural use despite presumed benefit of the clock acceleration. Delivery of specific RNP into barley calli mitotic cells by gold particle bombardment, followed by next-generation sequencing and droplet digital PCR (ddPCR) demonstrate a significant increase of recombinant alleles in treated vs control samples. These results are demonstrating that transient induction of specific DSB in mitotically-dividing cells can be exploited for transgene free and precise breeding of crops.

## Materials and Methods

### Plant material and genotyping

The HEB-25-049 plant material used in this experiment is originated from the barley NAM population HEB-25 (Maurer et al. 2015). This line with residual heterozygosity was further selfed to generate BC1S5 segregating seeds and plants for phenotype and RNP delivery experiments. A genotyping assay for the *Heat3.1* locus was developed using TaqMan®SNP assay. Each probe was designed to attach to a different nucleotide at in position 66,850,140 bp on chromosome 3H (A/T). The wild barley *Hordeum spontaneum* (Heat3.1^Hs^) probe was labeled with Yakima yellow dye and the cultivated *H. vulgare* (Heat3.1^Hv^) probe with FAM dye. We made 40X stock by combining 36µM and 8 µM of primer and probe, respectively, in a total volume of 100µl. We then used primers PM18.0020 and PM18.0021 (Supplementary Table 1) to amplify the desired region. Reaction mix for genotyping included: 10 ngr genomic DNA, 0.25 µl from the 40X stock in a 10 µl TaqMan®SNP assay mix. We made quantitative polymerase chain reaction (qPCR) steps as follows: Pre-PCR read 60°C, 30 sec; holding stage 95°C, 10min; 55 cycle of 95°C, 15sec; 60°C, 1min and finally a post-PCR 60°C, 30sec. Each assay included controls of plasmid or genomic DNA of known genotype (Heat3.1^Hs/Hs^; Heat3.1^Hv/Hv^; Heat3.1^Hs/Hv^).

### Field and circadian clock phenotyping

The field experiment was conducted in Agriculture Research Organization – Volcani center (ARO) in Rishon Lezion (E34º49’, N31º59’) in a nethouse during growing season 2017-2018. The nethouse is covered with a 50 mm MESH. Plants were sown into trays 171-hole planting trays containing soil mix (RAM 11), covered with Saran wrap and incubated for 48 hr at 4ºC to allow uniform germination, and moved to nethouse for further germination until transplanting (transplanting date January, 2017). Net house was divided into two parts: Eastern part that included fan was uncovered, while western part was covered with polyethylene, and three electric fan heaters (GL3-3000W) were installed to have heat stress effect. For planting we used two 6 × 0.3 m troughs (Mapal Horticulture Trough System, Merom Golan, Israel) filled with lower layer of 70% tuff medium and covered with Odem soil type (Toof Merom, Golan, Israel). Two-leaf stage seedlings were transplanted to both sides of each trough with 10 cm distance between plants. For each trough one irrigation dripping pipe (30cm between each drip points) was installed. In order to ensure no drought or limited nutrient stress developed, plants where water irrigated 3 times a day combined with NPK fertilizer. We scored the genotype of all individual plants using a TaqMan®SNP assay (see above) and plant phenotyping was conducted as described previously (Merchuk-Ovnat et al., 2018).

In the circadian clock experiments, 24-36 BC1S5 plants were sown in 6-pack plastic trays (Tivan Biotech #1206 and #1020) and moved to 4ºC for at least 24 for unified germination. The SensyPAM room is thermally controlled thus allowed us to conduct experiments in high temperatures (HT, 34ºC) and optimal temperatures (OT, 22ºC) under continuous light. We made experiments in two environments to calculate the plasticity of the clock or the delta between values at HT and OT(Bdolach et al., 2019). The experiments started under OT conditions for 3 days in SensyPAM. We then returned the plants for three days under long days (14 hr of light) for entrainment in the growth room, before continuation to HT experiment for three more days. The F measurements of leaf areas were used to calculate photosynthesis parameters including NPQlss 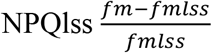 following (Bdolach et al., 2019). NPQlss was then used to calculate the free running period (FRP), amplitude (AMP), and relative amplitude error (RAE). The latter is a cumulative bias of the real measured values from the modelled cyclic line. These calculations made use of the BioDare2 package (https://biodare2.ed.ac.uk/) by setting the data input as “cubic dtr” and choosing the analysis method “mFourfit” (Zielinski et al., 2014).

### Statistical analysis

We performed the comparison between mean values under the different environments, and for the different traits, using the JMP14 software (SAS Institute Inc., Cary, NC, 1989-2019). After verifying normal distribution of values, we used one-way ANOVA to examine significant differences between the Heat3.1^Hs/Hs^; Heat3.1^Hv/Hv^; Heat3.1^Hs/Hv^ genotypes. For the circadian clock period, the delta between period of the same plants under high and optimal temperatures was compared between genotypes(Bdolach et al., 2019).

Since recombination rates among RNP-treated calli were not normally distributed, we compared these values between different treatments by using a non-parametric Kruskal-Wallis test (Ben-Israel et al., 2012).

### Cas9/gRNA in vitro assays with cloned PCR products as template DNA

We prepared two plasmids, PL18.0015 and PL.0018, which carry the Hv and Hs parental sequences by amplifying the target region with specific primers PM18.0020 and PM18.0021 (Supplementary Table 1) and using DNA that we extracted from Barke and HEB-25-25.1, respectively. After Sanger verification of these two plasmids we used them to amplify 5’ and 3’ of the 178bp piece with the primers PM18.0002 and PM18.0024 (5’ end), and PM18.0023 with PM18.0021 (3’ end; Supplementary Table 1). We then tailored these 5’ and 3’ amplicons by a second PCR with PM18.0020 and PM18.0021 to generate 4 possible combination (2 parental and two recombinants; Supplementary Table 2). The amplicons were extracted from a gel with PureLink™ Quick Gel Extraction Kit (Thermo, cat. no. K210012) and cloned to pJET1.2/blunt with CloneJET PCR Cloning Kit (Thermo, cat. no. K1231). The plasmids were transformed to Stellar™ Competent Cells (Invitrogen, cat. no. 636766) using a standard heat shock protocol (30 min on ice, followed by 45 sec at 42°C, adding 250 µL LB and shaking at 37°C for an hour), followed by selection on LB agar plate with ampicillin. The Plates were incubated at 37°C overnight and colonies were verified by colony PCR with pJET1.2/blunt primers. Another verification was done by digesting plasmids with restriction enzymes XhoI & BsaI (NEB). Finally, the plasmids were sequenced, and the right order of the SNPs was confirmed.

The synthetic sgRNA composed of universal tracrRNA and unique crRNA, were ordered from Invitrogen (cat. no. A35512&A35507). The synthetic crRNA and tracrRNA was resuspended with 1x TE buffer to 100µM. We conducted the PCR annealing of the crRNA and tracrRNA following the Invitrogen user guide protocol for a final crRNA:tracrRNA duplex concentration of 20 µM. We conducted *in vitro* assays following the BioLabs protocols. To obtain the best cleavage efficiency, the molar ratio of Cas9 sgRNA per target site was kept at 10:10:1 respectively. We included in a total volume of 24 µl reaction 3µl 10X Cas9 Nuclease Reaction Buffer, 3µl of 2000n (200nM final), 1µl of 6000nM *Streptococcus pyogenes* Cas9 Nuclease (New England Biolabs, cat no. M0386; 200nM final), and pre-incubate this for 10 min. at 25⁰C. We then added 6 µl of plasmid (20nM final), incubated the reaction for 15 min. more at 37°C, and stopped the digest by adding and mixing 1 ul of Proteinase K. The products and negative control (no sgRNA) were loaded on 1.5% agarose gel to test for linearization of plasmid.

### RNP preparation and particle gold preparation

We adopted protocols from (Barcelo and Lazzeri, 1995) and (Sparks and Jones, 2009) for the delivery of RNP and plasmids into calli using gold particles. We weighed ten mg of BIO-RAD sub-micron gold particles (0.6µm) and added 1ml of 100% ethanol within a 1.5ml Eppendorf tube. We then sonicated sample for two min and pelleted particles by centrifugation at 13,000 rpm for 10 to 30 sec, and discarded supernatant. We repeated this ethanol twice more with vortexing and tap mixing instead of sonication.

We then added 1ml of sterile water to the tube and sonicated sample again for 2min and pulsed spinned in a microfuge for 3 sec, followed by supernatant discard. Finally, we resuspended gold particles with 500µl of sterile water, divided into 50µl aliquots and stored tubes at −20°C. Assembly of the RNP was following Liang et al., 2018. For ten reactions of RNP delivery we prepared a total mix of 100 µl including 4µl 10X Cas9 protein (20 µg), 20µl of sgRNA(20 µg), 10 µg of 10X Cas9 reaction buffer, 66 µl RNase-free water and incubated this reaction at 25°C for 10 min. Prior to the use of golden particles we thawed the 50µl aliquots at room temperature and sonicated sample for 1-2 min. We then added this to the 100 ul RNP mix and gently yet thoroughly homogenized this by pipetting. We spread 15µl of the mixture onto the central region of each macro-carrier. The macro-carriers were air-dried on the benchtop at room temperature for 1h. The delivery of RNP into the calli cells that were positioned on callus induction media followed the book “Transgenic Crops of the World: Essential Protocols - Google Books,” n.d.. However, we changed the loading volume for each shoot to 15-20µl vs. 5 µl in the original protocol and set the rapture disk to 1100 psi, and positioned the target cells (plate) 7cm below the stopping screen. After delivery of the RNP into calli we kept the plates for 48 hours in dark before harvesting for DNA extraction as a pool or individually.

### Estimating recombination rate by Droplet Digital PCR (ddPCR)

The ddPCR reactions were conducted on the BioRAD QX200 Droplet Digital PCR System following manufacturer protocols (https://www.bio-rad.com/en-dz/product/qx200-droplet-digital-pcr-system?ID=MPOQQE4VY). Two primers and four distinct probes were designed; the first two probe are reporter fluorescent FAM and HEX attached to the first two SNPs of Heat3.1^Hs^ wild allele TTG: FAM attach to SNP1(**T**) and HEX to SNP2(**T**). In addition, two dark probes were also designed, to prevent attachment of FAM and HEX to SNP1 and SNP2 of Heat3.1^Hv^ cultivated allele **AC**A (Fig. 2). The Initial adjustments of the ddPCR reaction was making use of the four cloned plasmids that represent the four haplotypes (Supplementary Table 1). We made a serial dilution of the plasmid between 500 and 4 picogram/µl. We then mixed the plasmids to create three different assays: (1) **TT**G and **AC**A, (2) **TC**A and **AC**A, (3) **AT**G and **AC**A to generate the orange, blue and green clusters, respectively. It appeared that between 500-20 picogram/µl concentration, the plot result represented only a black cluster, considers all droplets as negatives. However, at 4 picograms, we could see the expected cluster distribution. Therefore, in the following ddPCR assays the plasmid diluted further to 12.5 femtogram/ul. Finally, we set channel one and two amplitude thresholds to 1051 and 2375, respectively, and used a concentration between 4 picograms/ul to 400 femtogram/ul since it was most suitable for plasmids detection with the expected distribution of the droplet clusters. For the ddPCR on the callus samples we measured DNA by nanodrop and used 10 ngr per reaction.

## Results and Discussion

### The barley Heat3.1 introgression and its effect on plant phenotype

As part of a study on the effects of natural allelic variation on circadian clock and phenotypic plasticity(Bdolach et al., 2019), we analyzed portion of the wild barley (*Hordeum vulgare*) HEB-25 population for field and circadian clock phenotypes using the SensyPAM platform. This analysis pointed to pleiotropic effects of a wild introgression, in the short arm of chromosome 3, on the clock period and reproductive traits (Fig. 1A). In follow-up experiment with segregating BC1S5 plants carriers of the wild allele introgression in this region (genotype Hs/Hs) were significantly linked (Student’s t-test; P<0.05) with increased acceleration and shortening of the clock compared to those plants (Fig. 1C). Within this genomic interval resides the barley ortholog of the Arabidopsis GIGANTEA gene (Fowler et al., 1999), which was shown in several plant species to modulate the circadian rhythm and its response to temperature changes(Edwards et al., 2005). Besides, on the same introgression are closely linked and complementary genes, Btr1 and Btr2, which control *rachis brittleness* in barley(Pourkheirandish et al., 2015) (Fig. 1A). Recessive mutations in any of these genes turn the fragile *rachis* (*brittle*) into a tough *rachis* phenotype (non-*brittle*) (Fernández-Calleja et al., 2019). Notably, within the HEB population, despite at least four rounds of recombination before this population was genotyped with iSELECT SNP markers (Maurer et al., 2015), we could not detect recombination events between the two loci. All carriers of the wild allele for the GIGANTEA gene are also free-shuttering types. This is implying linkage drag between the three loci from wild barley, which deemed maladaptive for breeding despite presumed benefit of the clock acceleration. We therefore decided to focus on these loci and utilize the genetic diversity between the cultivated *H. vulgare* (Hv) and its wild relative *H. spontaneum* (Hs) for development and implementation of RECAS9. The RECAS9 relies on CRISPR-Cas9 mediated DNA double-strand breaks (DSB)(Sadhu et al., 2016), which should allow directed homologous recombination, loss-of-heterozygosity, and finally, finer mapping of the causal DNA diversity underlying any quantitative or qualitative variation.

**Figure 1.**
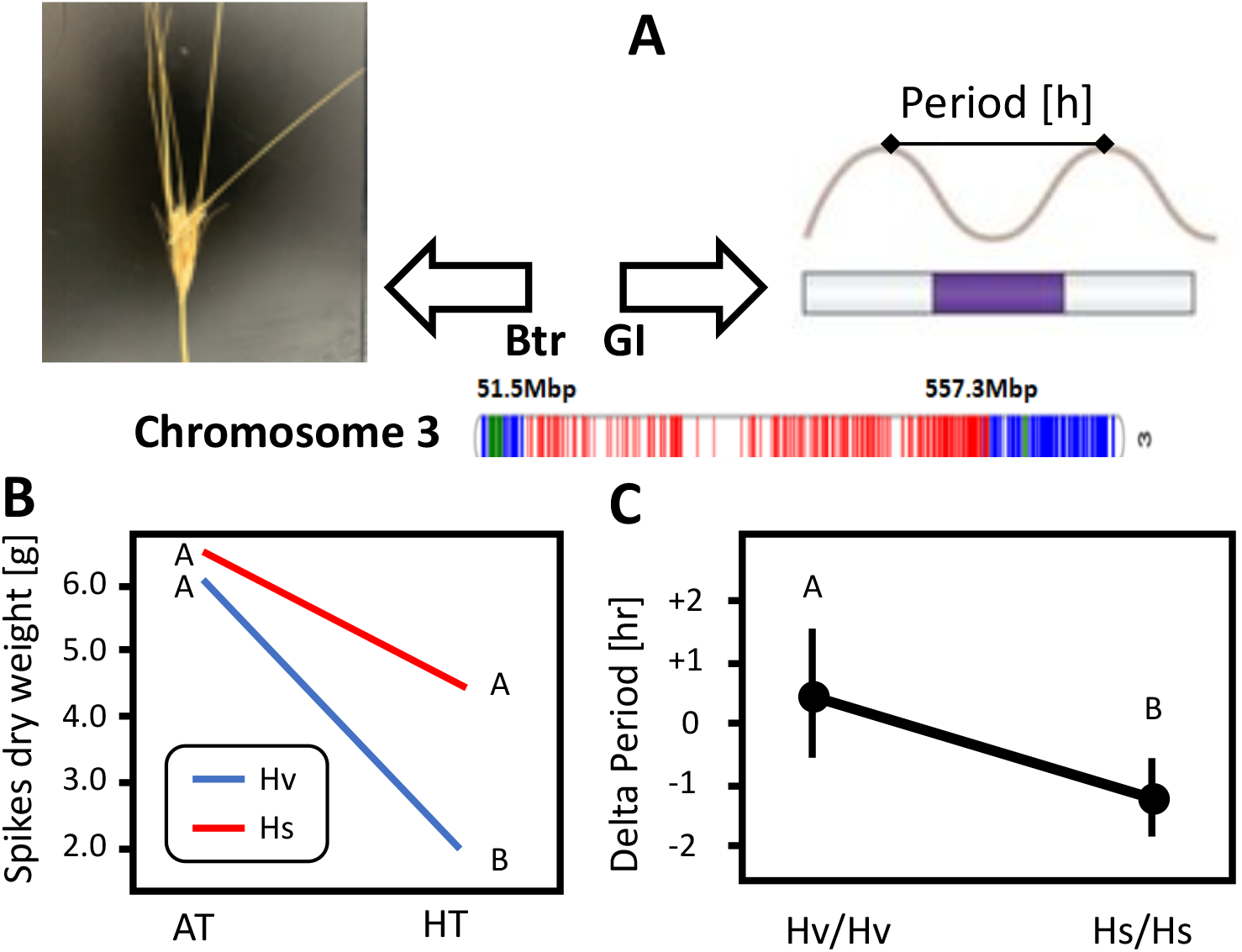
The genomic organization and phenotypic effects of the wild barley *Hordeum spontaneum* Heat3.1 locus. **A)** The wild introgression in line HEB-25-049, depicted in red, contains the Brittle rachis (Btr1 and Btr2) and GIGANTEA genes, which are involved in spike development and circadian clock rhythms, respectively. **B)** The *Heat3.1* locus is affecting the reproductive part under heat. Carriers of the wild (Hs) allele show less reduction of spike dry weight in transition from ambient to high temperature (AT to HT). **C)** Carriers of the wild allele show enhanced plasticity of the circadian clock rhythms under increased temperature. The delta in the clock period between high and optimal temperature is significant for carriers of the wild allele (Hs/Hs; mean acceleration of clock by 1.2 hr) as opposed to stable clock for those carrying the cultivated allele (Hv/Hv).

### In vitro assays of gene editing in the Heat3.1 locus

Our initial experiments included *in vitro* assays that tested the generation of DSB between SNPs by identifying 20 bp sequences upstream to protospacer-adjacent motives (PAM) in this region (Fig. 2A). We rationalized that the most effective way to test the efficiency, rather than specificity at this point, was to clone these targets within a cloning vector, and perform the *in vitro* assays (Fig. 1B). This plasmid linearization was deemed more clear than performing similar digests on PCR amplicons, since digestion product (linearized plasmid) and sgRNA leftovers did not overlap on gel. We obtained the gRNA by assembling specific crRNA and a general tracrRNA following manufacturer protocols (Invitrogen, see Methods). The 18.0083 and 18.0086 synthetic gRNA molecules were assembled with CAS9 protein to RNP, at the final molar ratio of 200nM(sgRNA):200nM(cas9):20nM(plasmid-template). We then incubated the RNP with the pJET plasmid, and both sgRNA molecules resulted in a specific digest and linearization of the plasmid (Fig. 1B).

**Figure 2.**
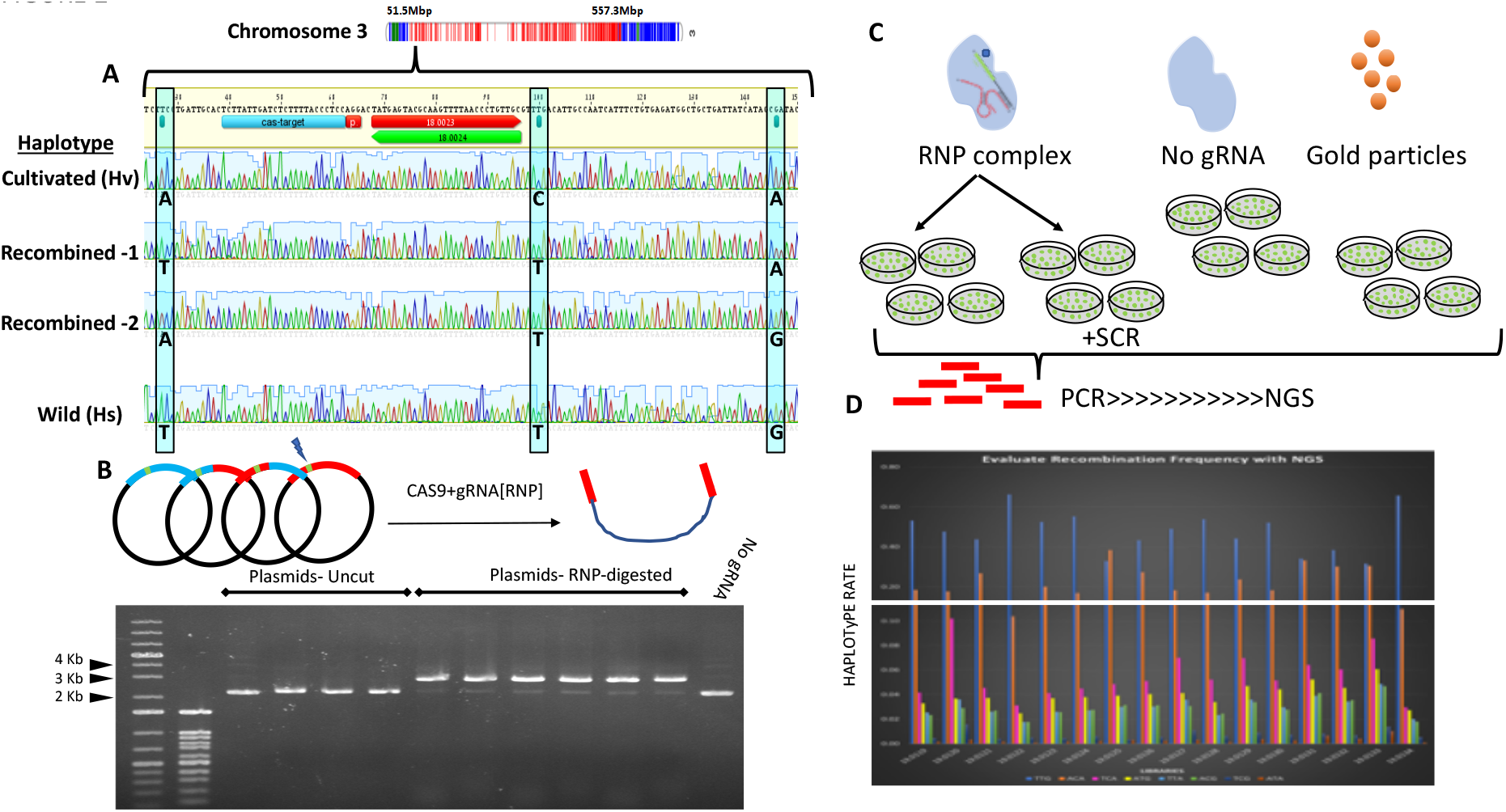
Haplotype organization, *in vitro* and *in vivo* assays for Cas9/sgRNA-mediated digestion and recombinogenic activity, respectively. **A)** The 178 bp including SNP 1 to 3 showing difference between Hs and Hv alleles. Above Sanger sequencing of the cloned amplicons indicated the Protospacer adjacent motif (*PAM*) in red and upstream sequence used as cRNA with tracrRNA to build the sgRNA. Downstream to these (green and red arrows) are the forward and reverse primers used to tailor the recombined alleles. **B)** The 178 bp were amplified and cloned into pJET plasmids and tested in vitro for specific digest. **C)** Calli of mixed homozygous and heterozygous calli were treated with RNP and controls, followed by PCR amplification of the 178 bp and **D)** Next generation sequencing analysis to determine the rate of the different haplotypes. Color coding for the parental (ACA and TTG) and other possible recombined haplotypes indicated below bar plot.

Before continuing from these *in vitro* assays to *in vivo* application of these RNP to plant mitotic cells, we have engineered similar plasmids that contains the two parental alleles, i.e. Hv and Hs (Fig. 1B). Besides, we recombined these allelic fragments by “tailoring PCR” (see Methods) and generated eventually four pJET plasmids containing a piece of 178 bp harboring the four haplotypes based on 3 SNPs (Hv, Hs and two recombinant products) (Fig. 2B). These plasmids served later as controls for measurements of recombination frequencies *in vivo*.

### In vivo assays of RECAS9: mitotic recombination mediated by RNP and analysis by next-generation sequencing

The RNP complex including the 18.0083 sgRNA located between the first and second SNPs, which we validated in the *in vitro* assay (Fig. 1C), delivered by gold particle bombardment into calli cells (see Methods). These calli were originated from mature embryos segregating for the *Heat3.1* locus (self of heterozygous plant in the BC1S5 generation (Maurer et al., 2015)). For the most parts, biolistic delivery followed Liang et al., 2018 with few modifications (See Methods). We divided the first experiment into six treatments (**I-VI**) that included three replicates of ten calli in each plate. All samples contained the two components of RNP complex Cas9 protein with sgRNA, except for the negative control (**VI**) which contained only the Cas9 protein. Besides, to the RNP complex groups **I** and **II** we performed co-bombardment of a plasmid containing ‘Helper morphogenic genes’ BABYBOOM WUSCHEL in the thought to achieve accelerated cell divisions and theoretically increased recombination rates(Svitashev et al., 2016) (Lowe et al., 2016). We added to the media of groups **II** and **IV** the SCR7 substance. The SCR7 is an inhibitor for the non-homologous end joining (NHEJ) process (Ma et al., 2016)(Chu et al., 2015)(Maruyama et al., 2015). Group **V** incubated with high osmotic media one day before the bombardment (Supplementary Fig. 1A). The bombardment followed by PCR-based library preparation for next-generation sequencing to estimate recombination rates. This experiment included eighteen libraries (six treatments × 3 replicates). DNA extraction and PCR amplification with a barcode-linker were conducted for each library to allow multiplexing. The amplicon libraries analyzed by MiSEQ sequencing using 150bp paired-end (PE150) (Supplementary Table 1). Besides, we included the four cloned plasmids with parental and recombined alleles as a control template for the PCR reactions and the NGS.

We analyzed NGS results for both HDR and NHEJ. As expected, the cloned plasmids showed only one dominant allele, with a low and neglected rate of the other alleles (Sup Fig. 1B). We identified the HDR products by scoring the different alleles and their frequency; TTG and ACA are the parental Heat3.1^Hs^ and Heat3.1^Hv^ alleles, respectively. The four main recombinant products are TCA, ATG, TTA, and ACG. TCA and ATG generated from DSB between SNP1 and SNP2 and TTA and ACG from DSB between SNP2 and SNP3. Besides, TCG, ATA correspond to double recombination, i.e. one between SNP1 and SNP2 and second between SNP2 and SNP3. We observed recombination in all the different treatments with a higher frequency of TCA and ATG recombinants. However, we could not identify a significant change between the treatments with regard to the proportion of the recombinant compared to the parental alleles (Sup Fig. 2B). Since the negative controls showed similar recombination rates as the other treatment, the bombardment repeated with more negative controls replicates, to refute, for example, the possibility that the negative control also contained the sgRNA as the examined samples.

In the second bombardment, we mainly wished to understand why we found recombination frequency also in the negative controls. This experiment included four treatments, and each treatment had four replicates, together we had sixteen plates with ten calli in each one. The RNP group divided into two, four plates with SCR7 and four plates without SCR7. Two negative control treatments included Cas9 protein without sgRNA or only gold particles (Fig. 2C). Again, analysis of the NGS results indicated that under this experimental set-up, where each sample composed of segregating calli (homozygous and heterozygous) recombination rate is on average 4.8%, yet not significantly different between RNP and negative controls (Fig 2B).

### In vivo assays of RECAS9: mitotic recombination mediated by RNP and analysis by digital droplet PCR (ddPCR)

One possible reason for the identification of the recombinant product in the negative controls, as well as in the RNP treated samples, is that recombination levels *in vivo* are within the range of recombination caused by PCR reactions. Meyerhans et al., 1990 estimated that the frequency of such recombination is up to 5.4%, and suggested that these recombinational events are a consequence of prematurely terminated products acting as primers by annealing to partially homologous templates during subsequent PCR cycles. Odelberg et al., 1995 exhibited additional mechanisms that can contribute to generating recombinant molecules as a template-switching process, in which the polymerase or the nascent strand switches from the original template to a secondary template during DNA synthesis. Besides, a well-known phenomenon in recent microbiome studies is a chimeric error that can be corrected in the bioinformatics processing workflow (Karstens et al., 2018). Notably, Odelberg et al., 1995 showed that one could reduce recombination products by attaching and separating the complementary template to streptavidin magnetic beads.

Realizing this chimeric issue resulting from PCR and its possible causes, we switched to digital droplet PCR (ddPCR) as the detection method. One underlying assumption of dPCR is that it includes random distribution of DNA molecules into the partitions and amplification of a single target in a partition (Whale et al., 2013). In our study, this would be either the parental or the recombinant alleles in single droplets. In these ddPCR assays, two probes are the reporter fluorescent FAM and HEX attached to the first two SNPs of Heat3.1^Hs^ *H. spontaneum* (Hs) allele TTG: the FAM attach to SNP1(**T**) and HEX to SNP2(**T**). Besides, we designed two dark probes to prevent attachment of FAM and HEX to SNP1 and SNP2 of Heat3.1^Hv^ cultivated allele **AC**A (Supplementary Fig. 2A). Once recombined, the two probes are changing from CIS to TRANS configuration (Supplementary Fig. 2B). Consequently, FAM and dark probe attach to **TC**A recombinant and HEX and dark probe attach to **AT**G recombinant (Supplementary Fig. 2B). In the ddPCR output, **TT**G and **AC**A allele together emit orange and black colour, respectively. Therefore, orange and black clusters indicate no recombinant events, while blue color indicates for **TC**A recombinant and green color depict the **AT**G recombinant (Supplementary Fig. 2C). To test specificity of the probes for the different allele product we used the pJET plasmids with the different allele products. Indeed, after several experiments we determined the optimal probes and DNA template concentration that detected the different alleles and their mixes (Fig. 3A-C; see Methods).

**Figure 3.**
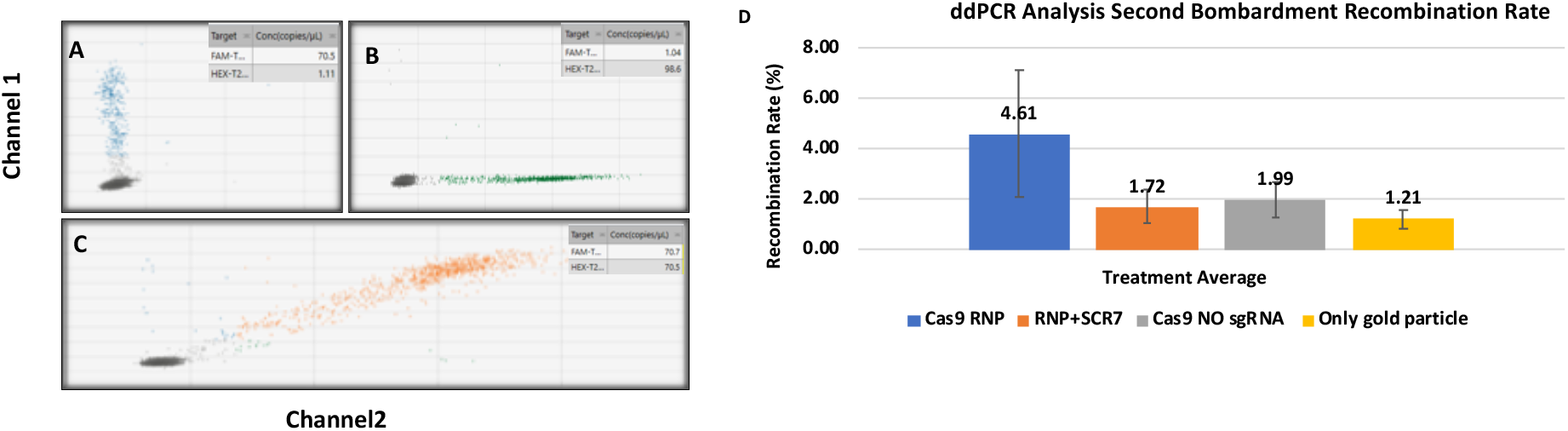
Droplet Digital PCR (ddPCR) assays for detecting and quantifying recombination levels in RNP treated calli. ddPCR using plasmids **A) TC**A& **AC**A templates **B) AT**G& **AC**A templates and **C) TT**G& **AC**A templates. Copies of target per microliter (copies/ul) are represent in the upper right insets. **D)** Mean percentage of recombination rates at the second bombardment. Y-axis is [(FAM-HEX)/HEX]*100 absulte value, indicating the recombination events with eror bars. Each treatmnet include four replicate with ten calli each.

In the following ddPCR experiments for the in vivo assays we estimated the recombination events by subtracting concentration of the FAM probe (copies/µL) from the HEX probe, and divided this delta by the smaller number. e.g. for sample 19.0120 this sum to 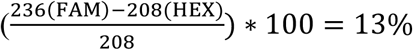 (Supplementary Table 2). These results showed that on average, calli undergoing Cas9 RNP treatment had higher recombination frequency than the other treatments (Fig. 3D). The negative results and particularly Cas9 without sgRNA treatment showed less than half of recombination frequency in the RNP treated calli (Fig. 3D). However, variation within treatments was relatively high, and differences between means were not significant according to parametric or non-parametric tests. We attempted to figure out the cause of the noise and hypothesized that one possible source could be the heterogeneity of the calli in a single DNA sample (10 calli each from a segregating batch of seeds; see Methods). We tested three sources of known genomic DNA from homozygote plants 16.0526 (Heat3.1^Hv/Hv^; ACA/ACA), 16.0527 (Heat3.1^Hs/Hs^; TTG/TTG) and Heterozygote 17-14e (Heat3.1^Hv/Hs^; TTG/ACA) in concentrations between 100ng to 1ng/µl. Supplementary Fig. 4 present the average of the eight dilution samples; results show unequivocally substantial noise in homozygote samples 16.0526 and 16.0527 with much higher noise for the samples of 16.0526 (CV=24%). As mentioned earlier, the dark probe attaches to **AC**A SNPs, and therefore 16.0526 samples should represent only within the black cluster. As opposed to the homozygous samples, the heterozygous genomic DNA presented much lesser variation between samples with most of the dilutions presenting no recombination as expected. Reason for this noise is since both probes are in CIS, and no extra FAM or HEX is expected (Supplementary Fig. 3).

Therefore, in the third bombardment, the experiment divided to four treatments, with each represented by six replicates (plates, with 20 calli in each): RNP complex, Cas9 expression cassette, Cas9 protein with no sgRNA, and only gold particle (Fig. 4A; see Methods). To avoid heterogeneity (mix of heterozygous and homozygous samples) and derived noise in ddPCR (see above) we scaled down DNA extraction methods. We used the TaqMan®SNP assay to distinguish between individual homozygote and heterozygous calli (Fig. 4B). We analyzed only heterozygote calli in the ddPCR for recombination events. Similar to the second bombardment (Fig. 3D), in this third bombardment with ddPCR analysis we identified higher recombination rates in the RNP treatment (4.77%) compared to the other treatment (Fig. 4D). Indeed, averaging of the different samples in each treatment showed a decreasing of the noise in all samples including negative controls (Cas9 without sgRNA and gold particle only) as reflected in the error bars (Fig 4C). Moreover, on average, there was a significant change in the average and distribution of recombination rate between RNP-treated to other treatments (Kruskal-Wallis test; P<0.0081), indicating that RECAS9 worked. Profoundly, at this point, there seem to be differential penetrance of the RECAS9 to the different samples, with some samples showing no recombination, while others are reaching 57% (sample 170; Fig. 4E). Attempts to improve the uniformity between samples by adjusting bombardment protocols, or alternatively with other delivery systems, are currently ongoing.

**Figure 4.**
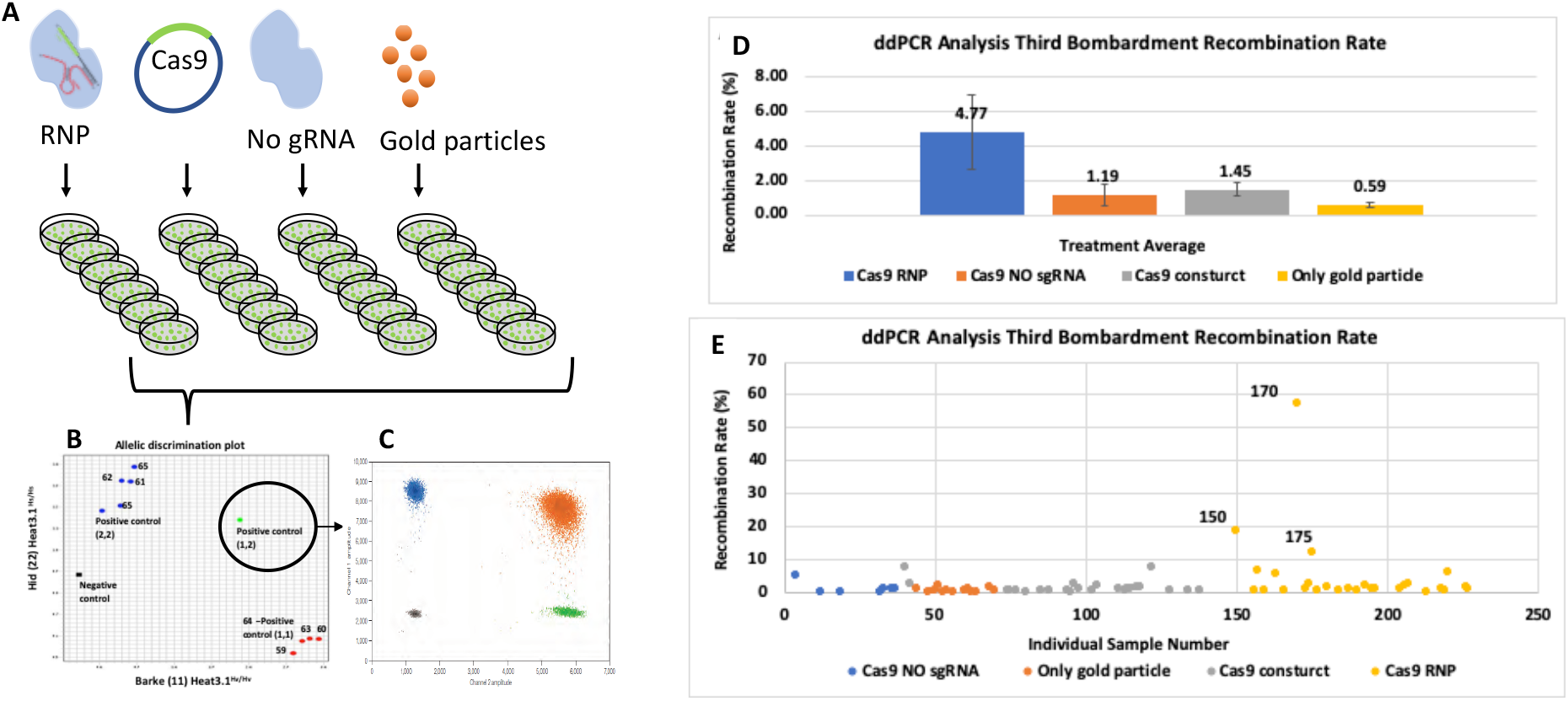
RNP delivered into calli heterozygous for the *Heat3.1* locus increases recombination rate. Calli originated from mature seeds segregating for the *Heat3.1* locus were **A)** treated with RNP or CAS/sgRNA plasmid contain the sgRNA between SNP1 and SNP2, followed by **B)** TaqMan®SNP assay to distinguish between the homozygotes to the heterozygotes, of which **C)** the later analyzed by ddPCR for recombination rates. **D)** A significant higher recombination rate in the Cas9 RNP (4.77%) treatment Y-axis is [FAM-HEX/HEX]*100 absulte value, indicating the recombination events with eror bars. Each treatmnet include four replicates with ten calli in each. **E)** Distribution of the recombination rate among individual calli samples as measured by ddPCR. Note the extreme high recombination rate found in several of the RNP-treated calli albeit the non-uniform penetrance of the treatment.

## Conclusions

The aim of this study is to allow high-resolution genetic mapping, and to break linkage drag, by increasing homologous direct repair (HDR) in mitotic cell (calli) following the Cas9-mediated double strand break (DSB). To our knowledge this is the first report of transgene-free, RNP mediated recombination, in plants. The study encounters a few challenges and surprisingly the main challenge is to estimate rare mutation events in heterogeneity callus cells since in low levels NGS resulted with confounding effects. Accelerating HDR events and regenerate a whole genetic modified plant, without selection marker rely on a precise and accurate detection assay. Nevertheless, the ddPCR method seems to overcome these difficulties although it does require extensive experimentation and DNA adjustments to make this a cost-effective tool, and for examining recombination beyond closely linked SNPs. Obviously, an efficient regeneration protocol from calli into plants would eventually allow screening and validating genetically modified plants by advanced allele mining approaches previously used in TILLING projects (Nida et al., 2016). Moreover, new Cas9 delivery systems, which have recently been developed like nanomaterials delivery(Demirer et al., 2019) and Haploid-Inducer Mediated Genome Editing (IMGE)(Wang et al., 2019)(Kelliher et al., 2019) can be utilized to enhance Cas9 delivery system and allow better penetration. On top of that, HDR process is less frequent in somatic cells then NHEJ, therefore it is requiring a procedure that allows increasing HDR over NHEJ pathway e.g. use specific inhibitors of NHEJ (as SCR7 in this study), or by engineering the Cas9 protein to fit better with mitotic recombination complexes.

## Acknowledgements

We wish to thank Tal Sherman (BioIlan, Givat Hen, Israel) for the assistance in setting up and applying the particle bombardment. We thank Eddy van Collenburg (Europe Bio-Rad Laboratories) for the guidance and assistance in setting-up the Droplet Digital PCR experiments. The IMOA Biotechnology grant #20-01-0136 to A.S and E.F supported this research. Work by M.R.P. was supported by the ARO postdoctorate fellowship.

## Contributions

S.L and E.F. designed the experiments, collected, analyzed and interpreted data, and wrote the manuscript. K.B. was in charge of the field experiments and the genetic analysis. M.R.P. was involved in the tissue culture activities, as well as in NGS and ddPCR analyses A.S. and E.F. supervised the genomic and genotypic data analysis.

---

**Supplementary Figure 1.**
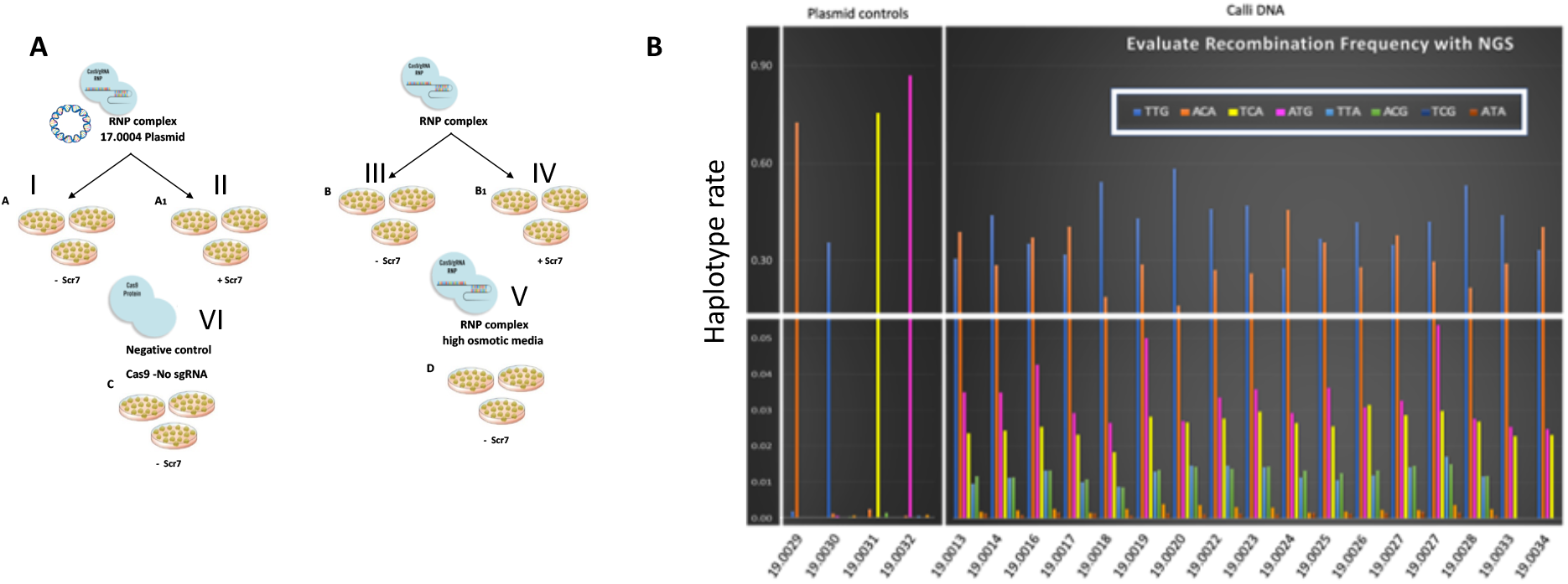
Next-generation sequencing for identifying recombination events in calli heterozygous and homozygous for the *Heat3.1*. **A)** Calli originated from mature seeds segregating for the *Heat3.1* locus were divided to five groups and treated as following: Groups I to V were treated with RNP complex. Groups I supplemented with a BBM WUS plasmid. In II and IV plates, media was supplemented with the NHEJ inhibitor SCR7. Group VI served as control to which we delivered Cas9 protein with no sgRNA. **B)** Next generation sequencing analysis for recombination rates. Bars are color-coded for the different 3 SNP haplotypes, including the parental TTG (Hs) and ACA (Hv) in blue and red colors, respectively. TCA, ATG, TTA, ACG represent the four main recombinant products. TCG and ATA represent double recombination scenario (Not in percent). On the left are the four PCR libraries from plasmids templates (19.0029-32), which include four cloned parental and recombined alleles. The other NGS libraries coding on the X-axis is detailed in Supplementary Table 1.

**Supplementary Figure 2.**
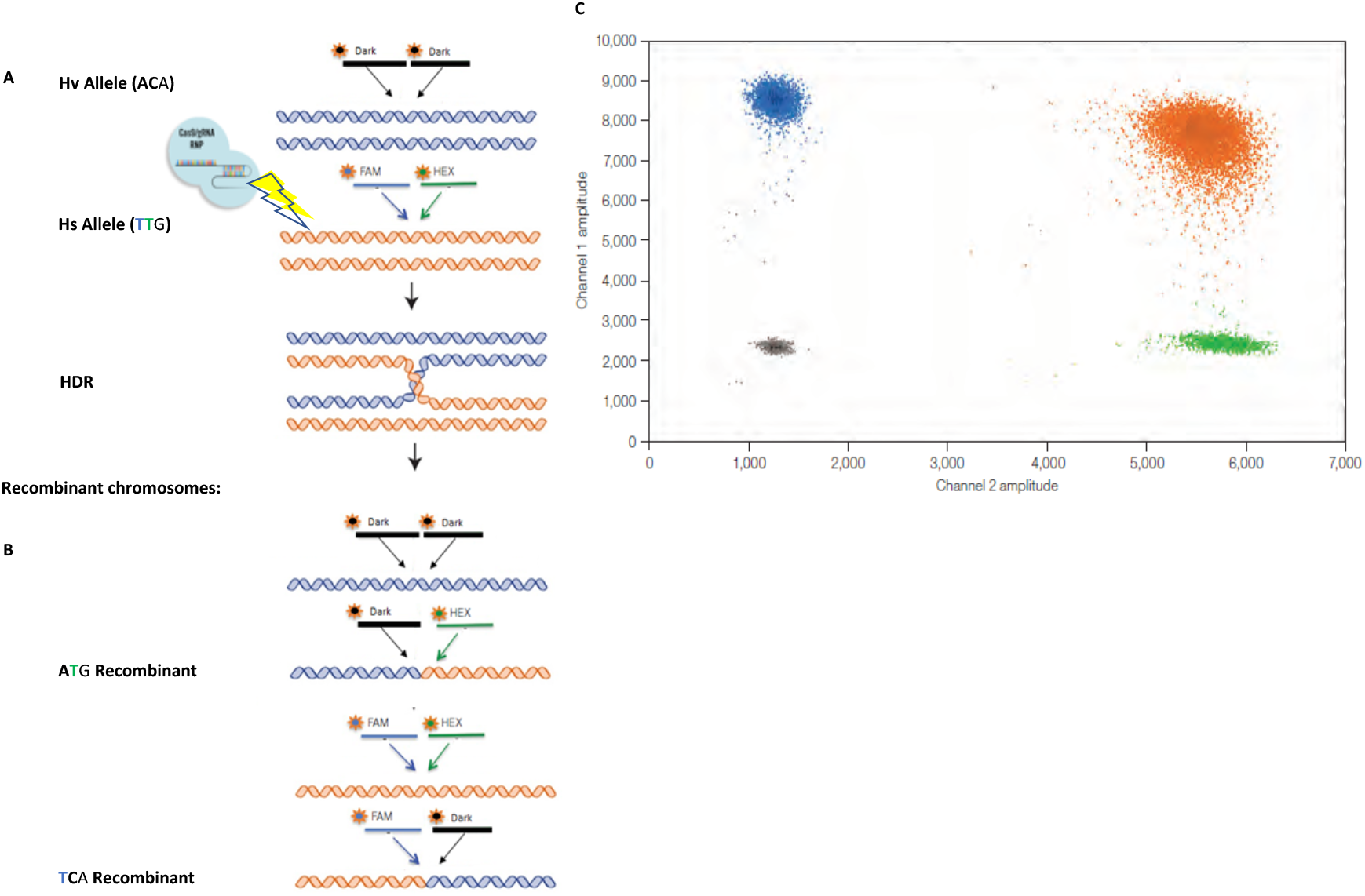
Design and principle of droplet digital PCR analysis assay for wild type and recombinant alleles. **A)** The FAM and HEX probe attach to the wild (Hs) SNPs, while two dark probes attach to the complementary cultivated (Hv) SNP **B)** DSB generated by RNP followed by HDR will generate two recombinant products. **B)** With no recombination, FAM and HEX probes attach to Hs allele and visualized only as orange cluster, while the dark probes attach to Hv allele and create black cluster. When some of the droplets includes recombinant template for the ddPCR, this FAM and dark probe, which attach to **TC**A recombinant, will generate blue cluster. Similarly, HEX and dark attach to **AT**G will generate green cluster.

**Sup FIGURE 3.**
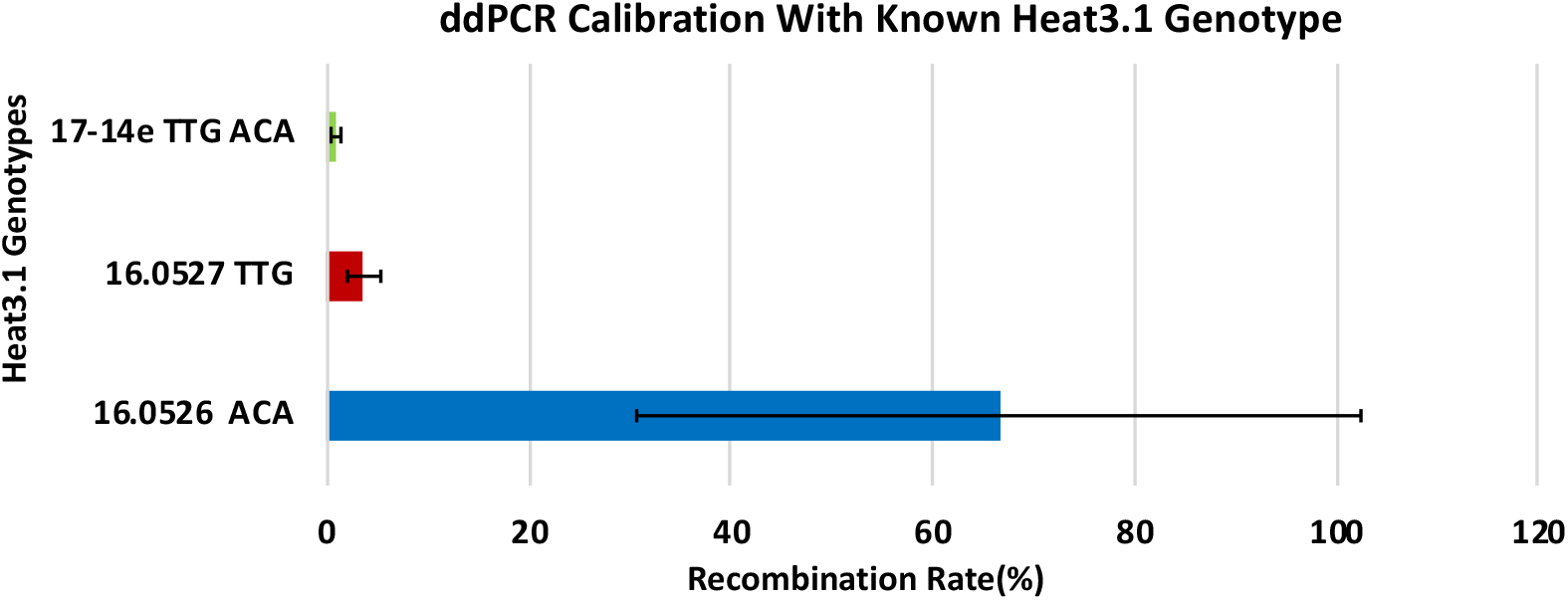

**Supplementary Table 1.**
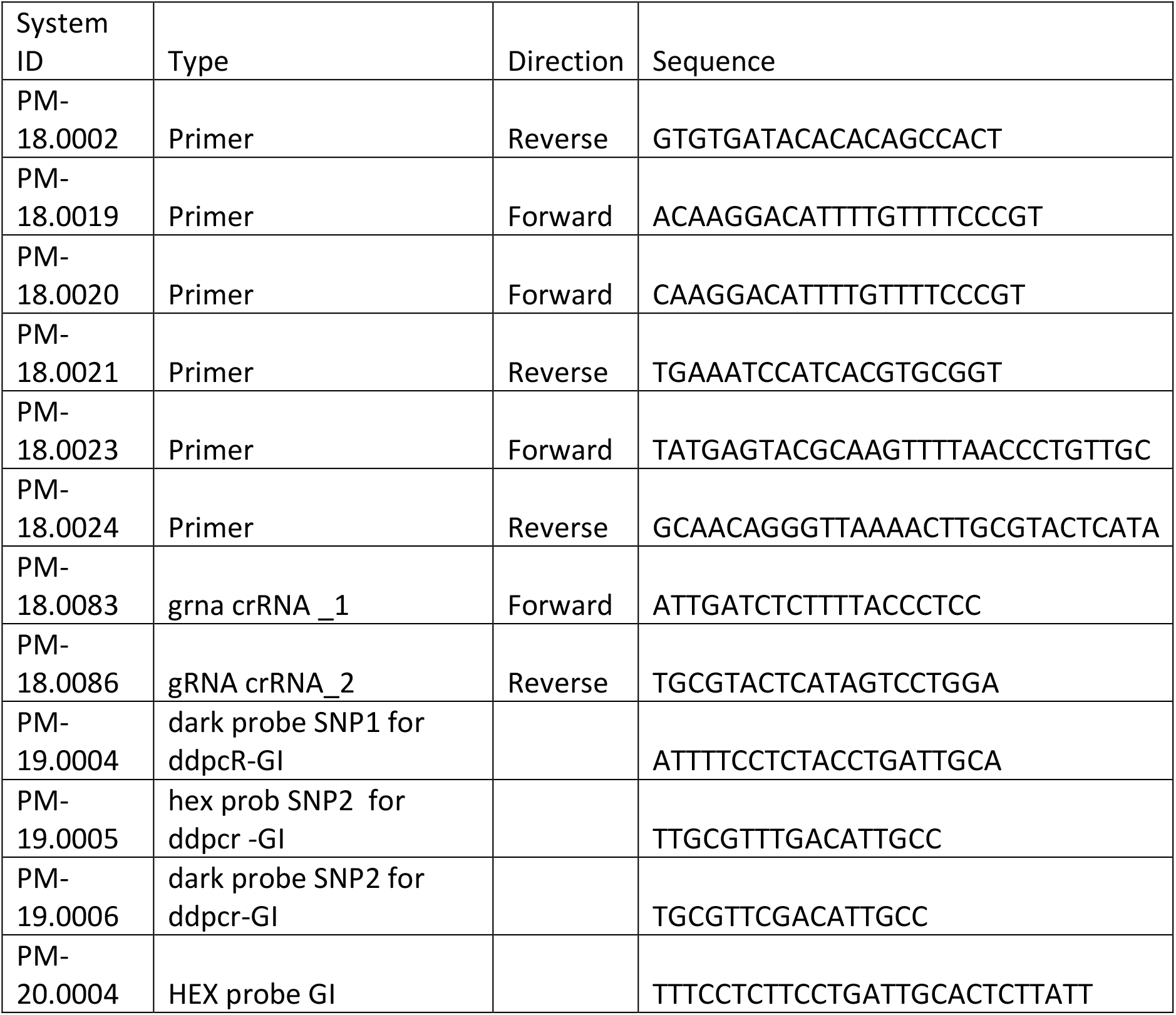
Primers and probes used in this study

**Supplementary Table 2.** Next generation sequencing libraries, first particle bombardment

| **System ID** | **Treatment** | **DNA template** |
| --- | --- | --- |
| SQ-19.0013 | II. RNP + BBM/WUS+Scr7 | Calli |
| SQ-19.0014 | II. RNP + BBM/WUS+Scr7 | Calli |
| SQ-19.0016 | II. RNP + BBM/WUS+Scr7 | Calli |
| SQ-19.0017 | I. RNP + BBM/WUS | Calli |
| SQ-19.0018 | I. RNP + BBM/WUS | Calli |
| SQ-19.0019 | I. RNP + BBM/WUS | Calli |
| SQ-19.0020 | IV. RNP + Scr7 | Calli |
| SQ-19.0021 | IV. RNP + Scr7 | Calli |
| SQ-19.0022 | IV. RNP + Scr7 | Calli |
| SQ-19.0023 | III. RNP | Calli |
| SQ-19.0024 | III. RNP | Calli |
| SQ-19.0025 | III. RNP | Calli |
| SQ-19.0026 | V. RNP + High Osmotic | Calli |
| SQ-19.0027 | V. RNP + High Osmotic | Calli |
| SQ-19.0028 | V. RNP + High Osmotic | Calli |
| SQ-19.0029 | pL18.0015 | Plasmid (ACA) |
| SQ-19.0030 | pL18.0018 | Plasmid (TTG) |
| SQ-19.0031 | PL.18.0017 | Plasmid (ATG) |
| SQ-19.0032 | PL.19.0005 | Plasmid (TCA) |
| SQ-19.0033 | VI. Negative control Cas9, No gRNA | Calli |
| SQ-19.0034 | VI. Negative control Cas9, No gRNA | Calli |

**Supplementary Table 3.** Next generation sequencing libraries, second particle bombardment

| System ID | Treatment |
| --- | --- |
| SQ-19.0119 | RNP-R1 |
| SQ-19.0120 | RNP-R2 |
| SQ-19.0121 | RNP-R3 |
| SQ-19.0122 | RNP-R4 |
| SQ-19.0123 | RNP + SCR7-R1 |
| SQ-19.0124 | RNP + SCR7-R2 |
| SQ-19.0125 | RNP + SCR7-R3 |
| SQ-19.0126 | RNP + SCR7-R4 |
| SQ-19.0127 | Negative control Cas9, No sgRNA- R1 |
| SQ-19.0128 | Negative control Cas9, No sgRNA-R2 |
| SQ-19.0129 | Negative control Cas9, No sgRNA -R3 |
| SQ-19.0130 | Negative control Cas9, No sgRNA -R4 |
| SQ-19.0131 | Negative control -only gold particle -R1 |
| SQ-19.0132 | Negative control -only gold particle -R2 |
| SQ-19.0133 | Negative control -only gold particle -R3 |
| SQ-19.0134 | Negative control -only gold particle -R4 |

